# Experimental infection of cattle with SARS-CoV-2

**DOI:** 10.1101/2020.08.25.254474

**Authors:** Lorenz Ulrich, Kerstin Wernike, Donata Hoffmann, Thomas C. Mettenleiter, Martin Beer

## Abstract

Six cattle (*Bos taurus*) were intranasally inoculated with SARS-CoV-2 and kept together with three naïve in-contact animals. Low-level virus replication and a specific sero-reactivity were observed in two inoculated animals, despite the presence of high antibody titers against a bovine betacoronavirus. The in-contact animals did not become infected.

---

After spill-over from a yet unknown animal host to humans, a global pandemic of an acute respiratory disease referred to as “coronavirus disease 2019 (COVID-19)” started in Wuhan, China, in December 2019 (1, 2). As causative agent, a novel coronavirus designated severe acute respiratory syndrome coronavirus 2 (SARS-CoV-2) was identified (3). Since the beginning of the pandemic, the role of livestock and wildlife species at the human-animal interface was discussed, with a special focus on the identification of susceptible species and potential reservoir or intermediate hosts. Until now, natural or experimental infections demonstrated the susceptibility of fruit bats (*Rousettus aegyptiacus*), ferrets, felids, dogs and minks, while pigs, chicken and ducks could not be infected (4–6). Besides ducks, chicken and pigs, major livestock species with close contact to humans are ruminants including a global population of ca. 1.5 Billion of cattle. In bovines, non-SARS betacoronaviruses are widespread (7, 8) with seroprevalences reaching up to 90% (9). The course of infection is usually subclinical (7). However, it is yet unknown whether any ruminant species including cattle is susceptible to SARS-CoV-2 infection or whether there is any cross-reactivity of antibodies against bovine coronaviruses (BCoV) to SARS-CoV-2.

## This study

To examine the susceptibility of cattle for SARS-CoV-2 and to characterize the course of infection under experimental conditions, six 4-5 months old, male Holstein-Friesian dairy calves were intranasally inoculated under BSL3-conditions with 1×10^5^ tissue culture infectious dose 50% (TCID_50_) of SARS-CoV-2 strain “2019_nCoV Muc-IMB-1” (GISAID ID_EPI_ISL_406862, designation “hCoV-19/Germany/BavPat1/2020”) at 1ml per nostril, using a vaporization device (Teleflex Medical, Germany). 24 hours after inoculation three contact cattle, that were separated prior to infection, were re-introduced. Body temperature and clinical signs were monitored daily and nasal, oral and rectal swabs were taken on days −1, 2, 3, 4, 6, 8, 12 and 20, and blood samples on days −1, 6, 12 and 20 after infection.

Swabs (Medical Wire & Equipment, UK) were immediately resuspended in 1.25ml serum-free cell culture medium supplemented with penicillin, streptomycin, gentamycin, and amphotericin B. Nucleic acid was extracted from 100μl of swab fluid using the NucleoMag Vet kit (Macherey-Nagel, Germany), and subsequently tested by the real-time RT-PCR “nCoV_IP4” targeting the RNA-dependent RNA polymerase (RdRp) gene (10). Positive results were confirmed by a second real-time RT-PCR based on an E gene target (11). Serum samples were tested by indirect immunofluorescence (iIFA) and virus neutralization assays (VNT) against SARS-CoV-2 as described before (5), and by an ELISA based on the receptor-binding domain (RBD) of SARS-CoV-2 (12). In addition, the sera were investigated by iIFA using CRFK cells (L0115, collection of cell lines in veterinary medicine (CCLV), Insel Riems) infected with BCoV strain Nebraska as antigen matrix and by VNT against this BCoV strain on MBDK cells (L0261, CCLV).

All animals tested negative for the presence of SARS-CoV-2 RNA in swab samples and SARS-CoV-2-specific antibodies in serum prior to infection. None of the inoculated cattle, nor any of the contact animals showed any clinical, disease-related symptoms. Body temperature, feed intake and general condition remained in a physiological range throughout the study. However, two of the inoculated animals became productively infected demonstrated by the detection of viral RNA in nasal swabs. One animal (number 776) tested positive on days 2 and 3 after inoculation with quantification cycle (Cq) values of 29.97 (day 2) and 33.79 (day 3), and another calf (number 768) on day 3 only (Cq 38.13) (Figure 1A). These animals scored positive only in the nasal swabs. Oral and rectal swabs taken simultaneously, as well as specimens collected from every other animal, remained negative throughout the study period.

**Figure 1.**
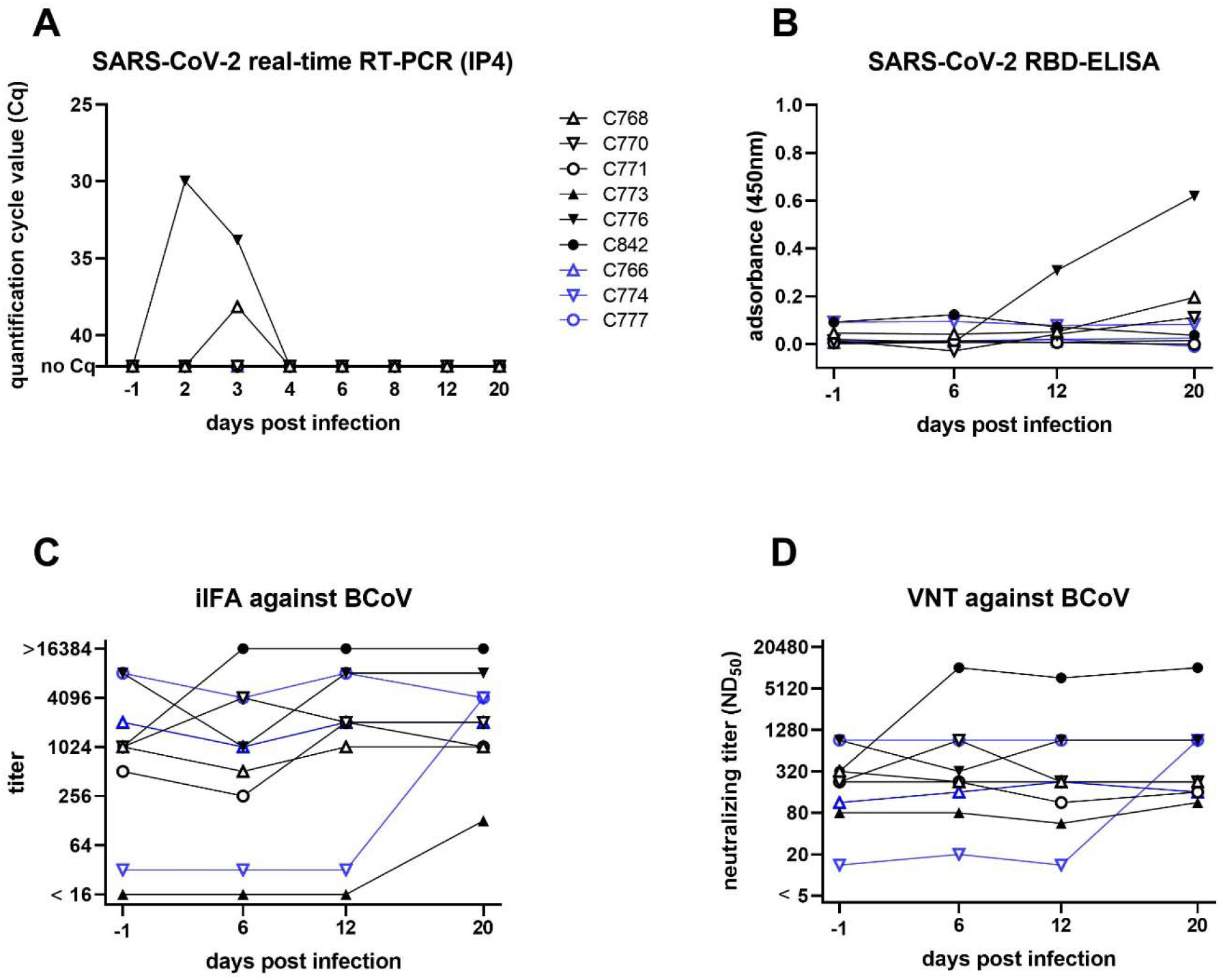
Characterization of a SARS-CoV-2 infection in cattle. Animals directly inoculated are shown in black, while in-contact animals are depicted in blue. Individual animals are indicated by the same symbol in every figure panel. (A) Viral load in nasal swabs measured by real-time RT-PCR. Cattle 776 and 768 presented with detectable viral loads in nasal swabs on day 2 and/or 3. (B) Results of an RBD-based SARS-CoV-2 ELISA for sera taken on days −1, 6, 12 and 20. (C+D) Serological status towards bovine coronavirus. Cattle 842, which tested positive for BCoV in the nasal swab by RT-PCR, presented with a titer increase in both indirect immunofluorescence (iIFA) (C) and virus neutralization test (VNT) (D). Pre-infection antibody titers against BCoV did not influence infection with SARS-CoV-2, as cattle 776 and 768, which tested positive for SARS-CoV-2 genome (panel A), showed no infection related reaction of BCoV antibody titers.

Serum samples were tested with a SARS-CoV-2 RBD-specific indirect ELISA. An increase in reactivity was observed for animal 776 from day 12 onwards (Figure 1B) indicating seroconversion. Serum taken on day 20 from this animal confirmed the positive ELISA findings with a low iIFA titer of 1:4, and a visible, although not complete, inhibition of viral replication in VNT (serum dilution 1:2). Animal 768 showed only a slightly increased ELISA-reactivity at day 20, while iIFA and VNT remained negative. This could be related to the test sensitivity or a possible restriction of replication to the upper respiratory tract.

The other animals remained negative throughout the study in all applied SARS-CoV-2-specific serological tests.

In addition, the BCoV status of the cattle was tested. As confirmed by VNT, all animals had neutralizing antibodies against BCoV prior to SARS-CoV-2 infection, but the titers differed markedly between individual animals (Figure 1D). Surprisingly, three animals showed an increase in antibody titers against BCoV in iIFA and two also in the VNT (Figure 1). In order to exclude an effect of the SARS-CoV-2 infection, nasal swabs were tested for BCoV by a generic RdRp-based RT-PCR (13). Animal 842 reacted positive one day prior to SARS-CoV-2 infection and 2 days post infection. The presence of a non-SARS-BCoV, which induced the increase in the anti-BCoV titer in this animal (Figure 1) and presumably infected animal 774, was confirmed by sequencing. However, no interference of the bovine coronavirus with the applied SARS-CoV-2 tests was observed, since all animals tested negative in SARS-CoV-2 tests prior to infection (Figure 1). Hence, there is presumably no detectable serological cross-reactivity between BCoV and SARS-CoV-2 in the used assays. Moreover, two animals with high BCoV sero-response were PCR-positive for SARS-CoV-2 RNA after inoculation, whereas those with lower BCoV-specific titers could not be infected, further confirming a lack of any cross-reactivity or cross-protection.

In conclusion, our findings demonstrate that under our experimental conditions cattle show low susceptibility to SARS-CoV-2, since two out of six animals appear to be infected as demonstrated by SARS-CoV-2-genome detection in nasal swabs and specific seroconversion. However, there is no indication that cattle play any role in the human pandemic nor are there reports of naturally infected bovines. This correlates with the rather low genome loads we detected after experimental intranasal infection of cattle and the absence of transmission to any of the direct in-contact animals. Nevertheless, in regions with high numbers of cattle and high case numbers in humans, like the US or South America, close contact between livestock and infected animal owners or caretakers could lead to anthropo-zoonotic infections of cattle, as it was already described for highly susceptible animal species like mink, felids or dogs (6, 14). Besides, age, husbandry practices and underlying health conditions of the animals should be considered, when assessing the risk of virus circulation within bovine populations. Hence, cattle may be included in outbreak investigations if there is any indication of direct contact to SARS-CoV-2, e.g. by infected farmers or staff. Beside direct detection by PCR, serological screenings with sensitive and specific ELISA-systems should also be taken into consideration. In this context, the wide distribution of another coronavirus in cattle is of special interest, especially since the presence of one virus did not protect from infection with another betacoronavirus in this study. Double infections of individual animals might potentially lead to recombination events between SARS-CoV-2 and BCoV, a phenomenon already described for other pandemic coronaviruses (15). A resulting chimeric virus, comprising characteristics of both primarily respiratory viruses, could present an additional threat for both human and livestock populations and should therefore be monitored.

## Acknowledgments

We thank Doreen Schulz, Bianka Hillmann, Mareen Lange and Constantin Klein for excellent technical assistance and the animal caretakers for their dedicated work. The study was supported by intramural funding of the German Federal Ministry of Food and Agriculture provided to the Friedrich-Loeffler-Institut and resources of the VetBioNet consortium (grant agreement No. EU731014), an initiative of the European Commission’s Horizon 2020 program.

## Ethical Statement

The experimental protocol was assessed and approved by the ethics committee of the State Office of Agriculture, Food Safety, and Fisheries in Mecklenburg-Western Pomerania (permission number MV/TSD/7221.3-2-010/18.

## References

1. WHO. COVID◻19 strategy update - 14 April 2020. Online available: https://wwwwhoint/publications-detail/covid-19-strategy-update---14-april-2020; last accessed: 16 May 2020. 2020.

2. Andersen KG, Rambaut A, Lipkin WI, Holmes EC, Garry RF. The proximal origin of SARS-CoV-2. Nature medicine. 2020 Apr;26(4):450–2.

3. Zhu N, Zhang D, Wang W, Li X, Yang B, Song J, et al. A Novel Coronavirus from Patients with Pneumonia in China, 2019. N Engl J Med. 2020 Feb 20;382(8):727–33.

4. Shi J, Wen Z, Zhong G, Yang H, Wang C, Huang B, et al. Susceptibility of ferrets, cats, dogs, and other domesticated animals to SARS–coronavirus 2. 2020;368(6494):1016–20.

5. Schlottau K, Rissmann M, Graaf A, Schön J, Sehl J, Wylezich C, et al. SARS-CoV-2 in fruit bats, ferrets, pigs, and chickens: an experimental transmission study. The Lancet Microbe. 2020.

6. Oreshkova N, Molenaar RJ, Vreman S, Harders F, Oude Munnink BB, Hakze-van der Honing RW, et al. SARS-CoV-2 infection in farmed minks, the Netherlands, April and May 2020. Euro Surveill. 2020 Jun;25(23).

7. Hodnik JJ, Ježek J, Starič J. Coronaviruses in cattle. Tropical animal health and production. 2020 Jul 17:1–8.

8. Tizard IR. Vaccination against coronaviruses in domestic animals. Vaccine. 2020 Jul 14;38(33):5123–30.

9. Boileau MJ, Kapil S. Bovine coronavirus associated syndromes. The Veterinary clinics of North America Food animal practice. 2010 Mar;26(1):123–46, table of contents.

10. WHO. Coronavirus disease (COVID-19) technical guidance: Laboratory testing for 2019-nCoV in humans. Online available: https://wwwwhoint/emergencies/diseases/novel-coronavirus-2019/technical-guidance/laboratory-guidance, last accessed: 16 May 2020. 2020.

11. Corman VM, Landt O, Kaiser M, Molenkamp R, Meijer A, Chu DKW, et al. Detection of 2019 novel coronavirus (2019-nCoV) by real-time RT-PCR. Euro Surveill. 2020 Jan;25(3).

12. Freuling CM, Breithaupt A, Mueller T, Sehl J, Balkema-Buschmann A, Rissmann M, et al. Susceptibility of raccoon dogs for experimental SARS-CoV-2 infection. bioRxiv : the preprint server for biology. 2020.

13. Dominguez SR, O'Shea TJ, Oko LM, Holmes KV. Detection of group 1 coronaviruses in bats in North America. Emerg Infect Dis. 2007 Sep;13(9):1295–300.

14. Sailleau C, Dumarest M, Vanhomwegen J, Delaplace M, Caro V, Kwasiborski A, et al. First detection and genome sequencing of SARS-CoV-2 in an infected cat in France. Transboundary and emerging diseases. 2020 Jun 5.

15. Forni D, Cagliani R, Clerici M, Sironi M. Molecular Evolution of Human Coronavirus Genomes. Trends in microbiology. 2017 Jan;25(1):35–48.

